# Passenger mutations in 2500 cancer genomes: Overall molecular functional impact and consequences

**DOI:** 10.1101/280446

**Authors:** Sushant Kumar, Jonathan Warrell, Shantao Li, Patrick D. McGillivray, William Meyerson, Leonidas Salichos, Arif Harmanci, Alexander Martinez-Fundichely, Calvin W.Y. Chan, Morten Muhlig Nielsen, Lucas Lochovsky, Yan Zhang, Xiaotong Li, Jakob Skou Pedersen, Carl Herrmann, Gad Getz, Ekta Khurana, Mark B. Gerstein

**Affiliations:** Program in Computational Biology and Bioinformatics, Yale University, New Haven, Connecticut, USA; Department of Molecular Biophysics and Biochemistry, Yale University, New Haven, Connecticut, USA; Department of Computer Science, Yale University, New Haven, Connecticut, USA; Yale School of Medicine, Yale University, New Haven, Connecticut, USA; Center for Precision Health, School of Biomedical Informatics, University of Texas Health Sciences Center, Houston, Texas, 77030, USA; Institute for Computational Biomedicine, Weill Cornell Medical College, New York, New York 10021 USA; Division of Theoretical Bioinformatics, German Cancer Research Center (DKFZ), 69120 Heidelberg, Germany; Faculty of Biosciences, Heidelberg University, 69120 Heidelberg, Germany; Institute of Pharmacy and Molecular Biotechnology, and Bioquant Center, University of Heidelberg, 69120 Heidelberg, Germany; Department of Molecular Medicine (MOMA), Aarhus University Hospital, Aarhus, Denmark; Department of Biomedical Informatics, College of Medicine, The Ohio State University, Columbus, Ohio 43210, USA; The Ohio State University Comprehensive Cancer Center (OSUCCC – James), Columbus, Ohio 43210, USA; The Broad Institute of MIT and Harvard, Cambridge, Massachusetts 02124, USA; Massachusetts General Hospital Center for Cancer Research, Charlestown, Massachusetts 02129, USA; Harvard Medical School, 250 Longwood Avenue, Boston, 02115, MA, USA; Department of Physiology and Biophysics, Weill Cornell Medicine, 1300 York Avenue, New York, NY, 10065, USA; Caryl and Israel Englander Institute for Precision Medicine, Weill Cornell Medicine, New York, NY, USA

## Abstract

The Pan-cancer Analysis of Whole Genomes (PCAWG) project provides an unprecedented opportunity to comprehensively characterize a vast set of uniformly annotated coding and non-coding mutations present in thousands of cancer genomes. Classical models of cancer progression posit that only a small number of these mutations strongly drive tumor progression and that the remaining ones (termed “putative passengers”) are inconsequential for tumorigenesis. In this study, we leveraged the comprehensive variant data from PCAWG to ascertain the molecular functional impact of each variant. The impact distribution of PCAWG mutations shows that, in addition to high- and low-impact mutations, there is a group of medium-impact putative passengers predicted to influence gene activity. Moreover, the predicted impact relates to the underlying mutational signature: different signatures confer divergent impact, differentially affecting distinct regulatory subsystems and gene categories. We also find that impact varies based on subclonal architecture (i.e., early vs. late mutations) and can be related to patient survival. Finally, we note that insufficient power due to limited cohort sizes precludes identification of weak drivers using standard recurrence-based approaches. To address this, we adapted an additive effects model derived from complex trait studies to show that aggregating the impact of putative passenger variants (i.e. including yet undetected weak drivers) provides significant predictability for cancer phenotypes beyond the PCAWG identified driver mutations (12.5% additive variance). Furthermore, this framework allowed us to estimate the frequency of potential weak driver mutations in the subset of PCAWG samples lacking well-characterized driver alterations.

## Introduction

Previous studies have focused on characterizing variants within coding regions of cancer genomes^1^. However, the extensive Pan-cancer Analysis of Whole Genomes (PCAWG) dataset^2^, which includes variant calls from more than 2500 uniformly processed whole-cancer genomes, offers an unparalleled opportunity to investigate the overall molecular functional impact of variants influencing both coding and non-coding genomic elements. Given that the majority of cancer variants lie in non-coding regions^3^, this variant dataset serves as a substantially more informative resource than the many existing datasets focused on exomes. Moreover, it also contains a full spectrum of variants, including somatic copy number alterations (SCNAs) and large structural variants (SVs), in addition to single-nucleotide variants (SNVs) and small insertions and deletions (INDELs).

Of the 44 million SNVs in the PCAWG variant data set, several thousand (less than 5/tumor^4^) are identified as drivers (i.e., positively selected variants that favor tumor growth) by recurrence-based driver detection methods. The remaining ∼99% of SNVs are termed passenger variants (referred to as *putative passengers* in this work), with poorly understood molecular consequences and fitness effects. Recent studies have proposed that, among putative passengers, some may weakly affect tumor cell fitness by promoting or inhibiting tumor growth. In prior studies, these variants have been described as “mini-drivers”^5^ and “deleterious passengers”^6^, respectively.

In this work, we explored the landscape of putative passengers in various cancer cohorts by leveraging the uniformly generated extensive pan-cancer variant calls^7,8^, driver mutation catalog^9^, transcriptome profile^10^, mutational signatures (see pcawg7 for details) and subclonal status^11^ of more than 2500 samples generated as part of PCAWG. Additionally, the driver discovery exercise^9,12^ in PCAWG suggests the absence of driver events in a few PCAWG samples despite having a large number of non-coding alterations. Thus, we closely inspected the cumulative effect of putative passengers in PCAWG samples. More specifically, we built upon and applied existing tools^13^ to annotate and score the predicted molecular functional impact of variants in the pan-cancer dataset. This systematic annotation and impact prediction effort generated a comprehensive compendium of PCAWG variants, thereby providing a valuable resource. Furthermore, we integrated the annotation and predicted impact score of each variant to quantify its overall impact on various genomic elements in different cancer cohorts. We observed that disruption of genetic regulatory elements in non-coding regions correlates with altered gene expression. Moreover, as elucidated by our signature analysis, various mutational processes have differential impact on coding genes and regulatory elements. We found that the predicted molecular functional impact of variants correlates with patient survival time and tumor clonality. We also observed differences in the predicted impact of putative passengers that may impact tumor progression. However, these putative passenger mutations may be driven purely by background processes or may suggest non-neutral effects. Hence, we also considered ways of assessing possible non-neutral roles for putative passengers.

We found that the aggregated impact of putative passengers provides significant predictive power (beyond common driver mutations) to distinguish cancer from non-cancer phenotypes, even after controlling for known mutational signatures and background mutation rates as possible confounders. We determined that this effect is prominent among tumors without known drivers, or with fewer driver variants than expected. Although the effects of these possible driver variants can only be detected in aggregate by our model, it motivates future searches for these variants among putative passengers, especially within non-coding regions of the genome. Finally, we note that selection acting on somatic cells is dynamic in nature. Thus, putative passengers may act as driver mutations at some later phase during tumor progression, when treatment is given, or when a clone spread to another organ. Therefore, it is valuable to characterize the functional impact of putative passengers, even when their functional effect is not of immediate selective consequence.

## Overall molecular functional impact

In order to characterize the landscape of putative passenger mutations in PCAWG (non-driver mutations based on the PCAWG driver catalog^9,12^), we first surveyed the predicted molecular functional impact (quantified by FunSeq score^13^) of somatic variants in different cancer genomes. The predicted functional impact distribution varies among cancer types and among genomic elements. A closer inspection of the pan-cancer impact score distribution for non-coding variants demonstrated three distinct regions. The upper and the lower extremes of this distribution are presumably enriched with high-impact strong drivers and low-impact neutral passengers, respectively. In contrast, the middle range of this distribution corresponds to putative passengers with intermediate molecular functional impact (**Fig 1a** & **supplement Fig. S6**).

**Figure 1:**
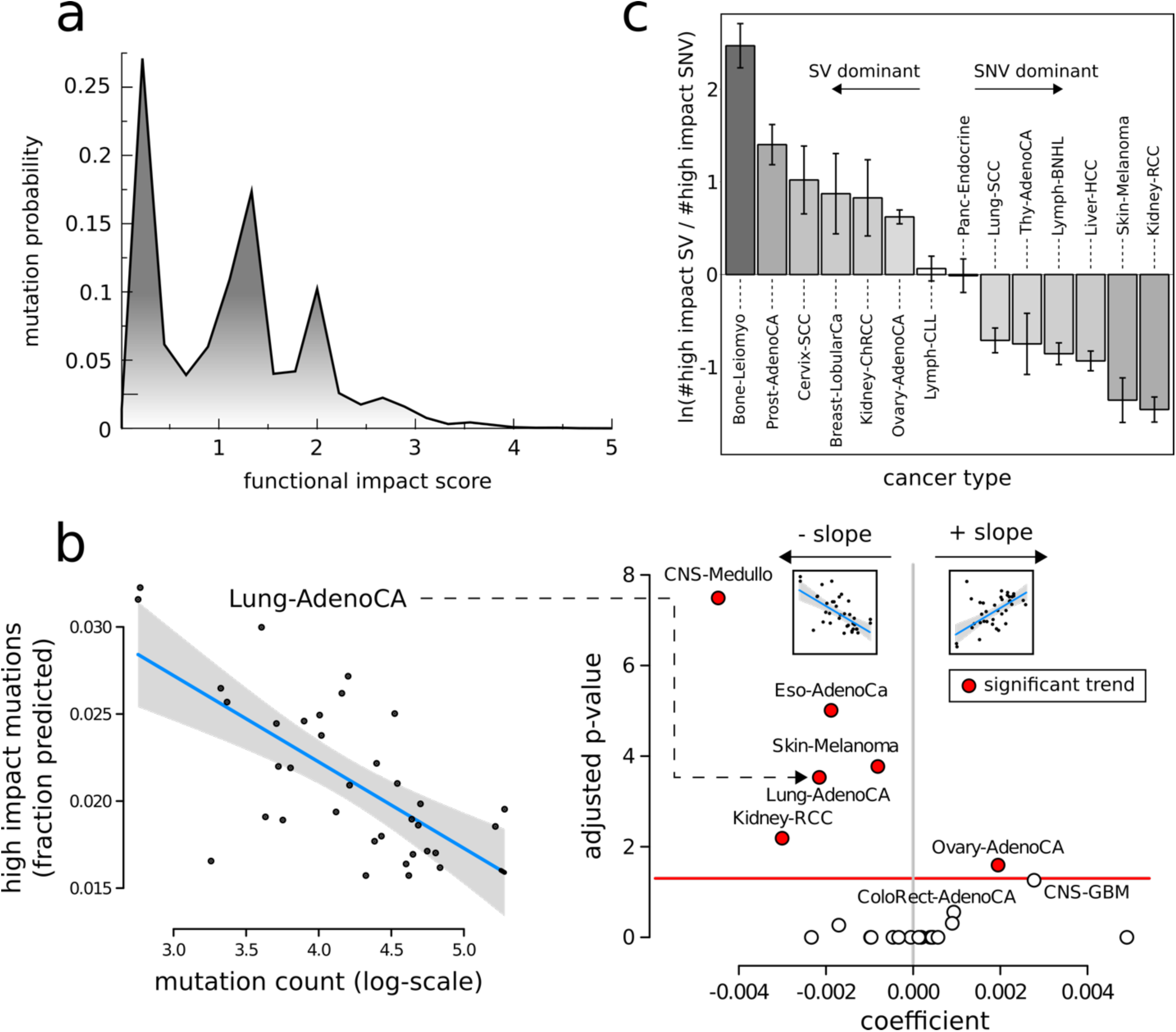
Overall functional impact of PCAWG variants. **a)** Functional impact distribution in non-coding(DHS) regions: three peaks correspond to low-, medium-, and high-impact variants;**b)** Correlation between the fraction of impactful non-coding SNVs and the total mutational counts for different cancer cohorts. **c)** log ratio of high-impact structural variants(SVs) and SNVs in different cancer cohorts. Error bar corresponds to variation among tumors within cohort.

Subsequently, we investigated whether the frequency of medium-and high*-*impact non-coding putative passengers (see **supplement section 1.4** for classification threshold) in a cancer cohort is proportional to its total mutational burden. For a uniform mutation distribution, we expect that the fraction of putative passengers would remain constant as cancer samples accumulate more mutations. In contrast, we observed that tumors with high SNV frequencies tend to have lower fraction of medium- and high-impact putative passengers. This trend is particularly strong in CNS medulloblastoma (p < 4e-8), lung adenocarcinoma (p < 3e-4), and a few other cancer cohorts (**Fig 1b** & **supplement Fig. S4-S5**).

In addition to SNVs, large structural variations (SVs) also play an important role in cancer progression. Thus, we quantified the putative functional impact of SVs (deletions and duplications). Briefly, we built a machine learning framework, which utilizes conservation, epigenomic signals and overlap with known cancer genes to assign a SV impact score (**supplement section 4.2**). A close inspection of both SV and SNV impact scores suggests that certain cancer subtypes tend to harbor many high-impact SVs, while others contain a large number of high-impact SNVs (**Fig 1c**). Many of these correlations have previously been observed^14^. For example, it is known that large deletions (on chromosome 17 in TP53 and BRCA1) play the role of drivers in ovarian cancer^14^, whereas clear cell kidney cancer is often driven by SNVs. However, we also find new associations, such as the predominance of high-impact large deletions compared to impactful SNVs in the bone leiomyoma cohort. Similarly, a close inspection of high-impact large duplications and high-impact SNVs suggests their differential affinity toward different cancer cohorts (see **supplement Fig. S7**).

## Burdening of different genomic elements

We investigated the overall mutational burden observed among different genomic elements in various cancer cohorts. Naively, one might assume that the overall burden of putative passengers in a cancer genome would be uniformly distributed across different functional elements and among different gene categories. In contrast, we observed that the predicted molecular impact burden in certain cancers is concentrated in particular regulatory regions and gene categories. This is easiest to understand in terms of coding loss-of-function variants (LoFs), where the putative molecular impact is most intuitive. We thus examined the fraction of deleterious LoFs affecting genes across several categories of cancer-related functional annotation (**Fig 2a** & **supplement Fig. S8-S9**). Driver LoF variants (included in the PCAWG driver catalog^9,12^) showed significant overlap with different categories of cancer-related genes (DNA repair, immune response, metabolic, and essential genes) relative to a uniform genome-wide expectation (p < 0.001). Conversely, non-driver LoFs displayed a small but significant depletion relative to a uniform genome-wide expectation in each of these categories except in metabolic and immune response genes, for which they showed slight enrichment compared to the genome-wide expectation (p < 0.001). We note that differential tendency towards mutation generation or mutation repair among these gene categories may contribute to these observations (e.g. higher expression among essential genes may lead to both increased transcription-coupled damage and transcription-coupled repair^15^).

**Figure 2:**
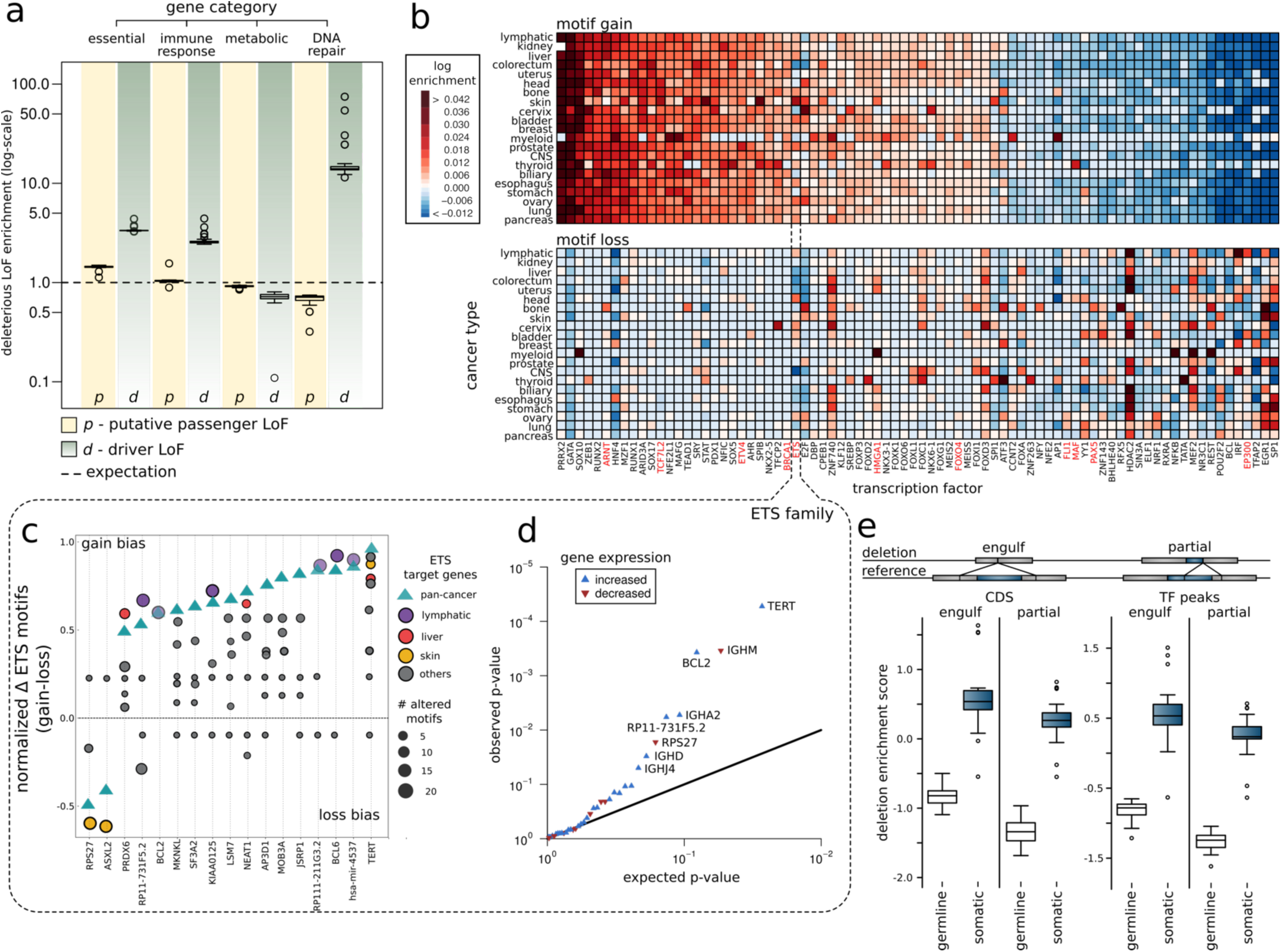
Overall functional burdening of different genomic elements. **a)** Percentage of genes in different gene categories (essential, immune response, metabolic and DNA repair genes) affected by driver and non-driver LoFs compared to uniform background expectation. Data points in boxplot corresponds to different tumor types.; **b)** Pan-cancer overview of TF burdening: heatmap of differential burdening of various TFs due to SNVs that induce motif-break and motif-gain events in different cohorts compared to a uniform genomic background; **c)** Target genes affected due to motif gain and loss in the ETS transcription factor family: genes such as TERT, RP17-731F5.2 and JSRP1 are affected due to gain-of-motif events, whereas ASXL2 and RPS27 are affected due to loss-of-motif events; **d)** Q-Q plot showing genes such as TERT, PIM1 and BCL2 that are differentially expressed due to gain-of-motif events in ETS TFs;**e)** Enrichment of germline and somatic large deletions in coding regions and transcription factor binding peaks. Large deletions can engulf or partially delete various genomic elements.

As with LoF variants, we can also quantify the overall burden of the non-coding SNVs in a cancer genome. However, for the majority of non-coding SNVs, predicted molecular functional impact is less easy to gauge. For instance, coding and non-coding variants occupying the terminal region of the gene or intronic regions would most likely have little functional consequence. In contrast, the molecular functional impact of transcription factor binding site (TFBS) variants is clearly manifested through the creation or destruction of transcription factor (TF) binding motifs (gain or loss of motif). In both cases (gain or loss), we observed significant differential burdening of TFBSs among different cancer cohorts. For instance, based on a uniform background model, we detected significant enrichment of variants creating new motifs in various TFs including GATA, PRRX2 and SOX10 (**Fig 2b**, **supplement table S12 & supplement Fig. S10-S11**) across major cancer types, compared with genome-wide expectation. Similarly, variants breaking motifs were highly enriched in TFs such as IRF, POU2F2, NR3C1and STAT in the majority of cohorts (**Fig 2b, supplement table S12** & **supplement Fig. S10-S11**). This selective enrichment or depletion suggests distinct alteration profiles associated with different components of regulatory networks in various cancers.

Furthermore, for a particular TF family, one can identify the associated target genes affected due to the bias towards creation or disruption of specific motifs in their regulatory elements (promoters and enhancers). For instance, the TERT gene shows the largest alteration bias for motif creation events among TFs in the ETS family across a variety of cancer types (**Fig 2c** & **supplement Fig. S12**). Other genes (such as BCL6) showed a similar bias, albeit in fewer cancers. Moreover, the enrichment of SNVs in select TF motifs leads to gain and break events in promoters that significantly perturb the overall downstream gene expression (**Fig 2d** & **supplemen**t **Fig. S12-S13**). For example, ETS family TFs at the regulatory region of TERT and BCL2 gene displayed a strong motif creation bias and a significant change in gene expression (with p-value TERT=5.49e-5 and p-value BCL2=3.4e-4). In contrast, RPS27 shows an alteration bias towards disruption of ETS motifs driven by skin melanoma, which coincides with a significant down-regulation of RPS27. Similarly, upon aggregating expression of all downstream genes affected by the same TF motif change, we found several motif alteration events significantly perturb downstream gene expressions. For instance, motif gain in ZBTB14 and E2F in lung adenocarcinoma, ETS motif gain in thyroid adenocarcinoma and TFAP2E and TFAP motif loss in lymphoma (**supplement Fig. S14-S15**).

Finally, we also analyzed the overall burden of structural variants (SVs) in various genomic elements and compared the pattern of somatic SV (large deletions and duplications) enrichment in cancer genomes to those from the germline (**Fig 2e**). As expected, we observed that somatic SVs were more enriched among functional regions compared to germline SVs, because the latter will be under negative selection for disrupting functional regions (**Fig 2e** & **supplement Fig. S16**). Furthermore, we observed a distinct pattern of enrichment for SVs that split a functional element versus those that engulf it. As has been previously noted, there is a greater enrichment of germline SVs that engulf an entire functional element rather than for those that break a functional element partially^16^. Interestingly, we observed the same pattern among somatic SVs (**Fig 2e** & **supplement Fig. S16**).

## Mutational processes analysis

The differential burdening of various genomic elements may be attributed to an underlying stochastic but uneven mutational processes. Thus, we closely inspected the underlying mutational processes generating SNVs in both coding and non-coding regions of cancer genomes. First, we looked into the most impactful event, loss-of-function mutations in coding regions. We would anticipate premature stop codons to show strong mutational contextual bias due to the nature of codon composition. For example, some mutations (e.g. T>Cs) do not result in a premature stop in any trinucleotide context. As anticipated, we found premature stop mutations carry a specific mutational spectrum, which differs significantly from the overall tumor mutational spectrum. Moreover, when compared with the pan-cancer premature stops, individual cancers display a spectrum shift. For example, premature stops in RCCs (renal cell carcinomas) show a higher percentage of T>As compared to all cancers (18% versus 8%) (**Fig 3a**). Similarly, a close inspection of penta-nucleotide spectrum associated with potential premature stop suggests high correlation between in-frame and out-of-frame mutational stop patterns (**supplement Fig. S17**). Moreover, the estimated ratio between observed and expected mutations suggests that Colorectal adenocarcinoma samples tolerate a greater number of loss of function mutations compared to Kidney-RCC and skin-melanoma cancers (**supplement Fig. S17**). Our observations can be explained by the divergence of mutational processes in individual cancer types and implies mutational processes confer distinct effects in coding regions.

**Figure 3.**
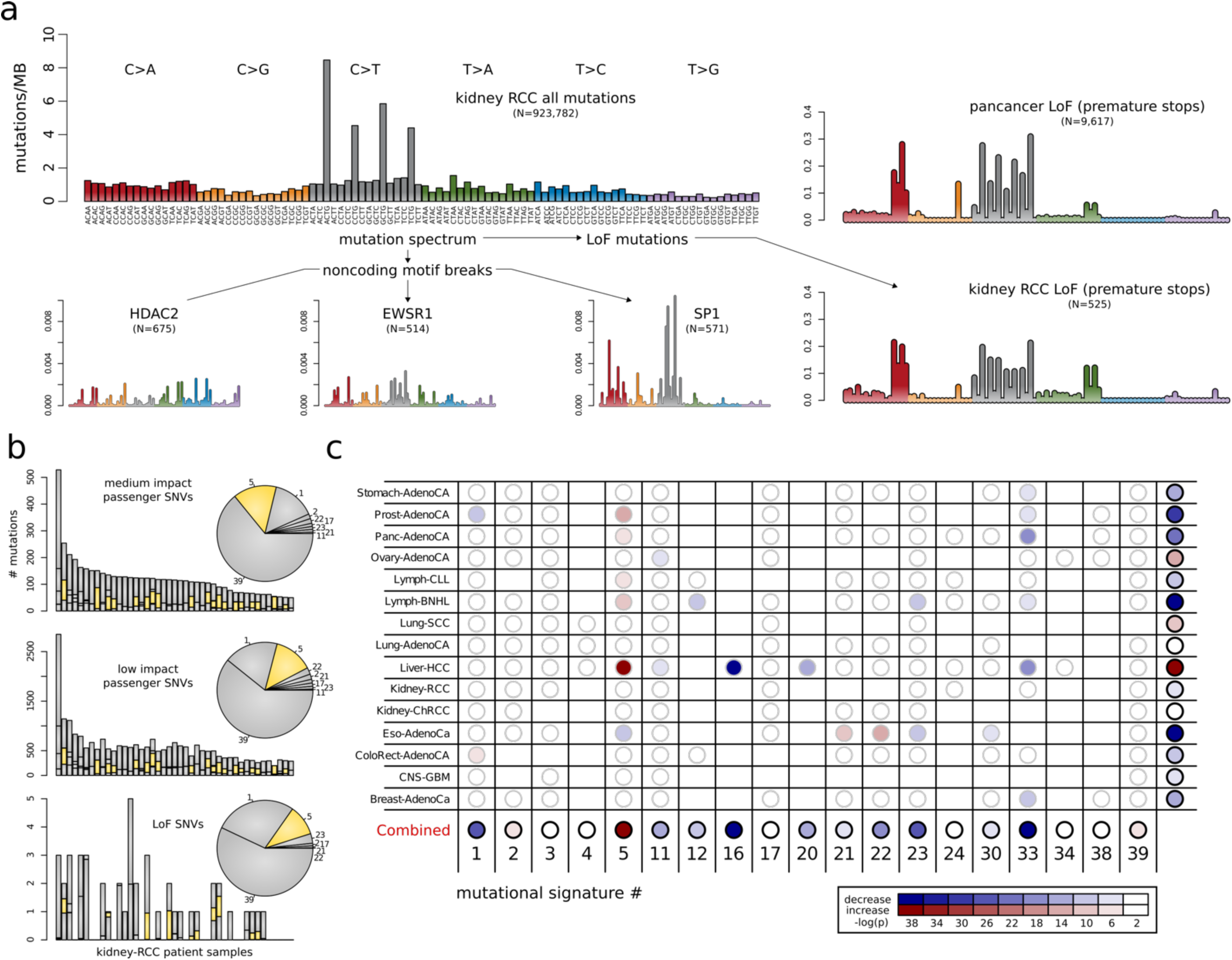
Mutational signatures associated with different categories of impactful variants. **a)** Mutation spectra associated with premature stops and transcription factor binding motif-breaking events observed in HDAC2, EWSR1 and SP1 in the kidney-RCC cohort. A pan-cancer premature stop spectrum is also shown for comparison; **b)** Distribution of mutational signatures in the kidney-chRCC cohort for impactful non-coding SNVs (top), low-impact non-coding SNVs (middle), and premature stops (bottom); **c)** Comparison of underlying signature distribution between high- and low-impact putative passengers in different cancer cohorts.

Similarly, the disproportionate functional load on certain TFs in cancers can be related to the underlying mutational spectrum influencing their binding sites. Different transcription factors have varying nucleotide context in their binding sites (TFBS). These variations may facilitate the role of certain mutational processes, which will be reflected in their mutational spectrum. For instance, the mutational spectrum of motif breaking events observed in SP1 TFBS suggests a major contribution from C>T and C>A mutation (**Fig 3a**). In contrast, motif-breaking events at the TFBS of HDAC2 and EWSR1 have relatively uniform mutational spectrum profiles.

Based on the mutational context, we can further decompose all observed mutations into a linear combination of mutational signatures, which presumably represent the mutational processes^17,18^. Every signature (see pcawg7 paper for details) has varying influence depending on the cancer type and in a given cancer type, different signatures disproportionally burden the genome. Comparing the signature composition of low- and high-impact putative passengers in certain cancer-cohorts can help us to distinguish between mutational processes that generate distinct variant impact classes. For instance, in the chRCC (chromophobe renal cell carcinoma) cohort, although the majority of putative passenger variants can be explained by signature 39, high-impact and low-impact putative passengers have a different proportion of signature 5 and signature 1(**Fig 3b**). We also scrutinized LoFs in coding regions, which presumably carry the highest molecular function impact. Compared to non-coding putative passengers, signature 1 and 23 together contribute a relatively higher fraction of premature stops. Similar to the Kidney-chRCC cohort, we observed distinct signature distributions for the low-and high-impact non-coding putative passengers in Liver-HCC, Prost-AdenoCA, Eso-AdenoCA and Ovary-AdenoCA cohorts (**Fig 3c**). In addition to mutational signatures, we observed that cancer samples with microsatellite instability (MSI) due to failure of DNA mismatch repair, have a higher percentage of high impact non-coding putative passengers (**supplement Fig. S18**). Collectively, these findings suggest that various mutational processes shape and disproportionally burden cancer genomes.

## Subclonal architecture and cancer progression

Cancer is an evolutionary process, often characterized by the presence of different sub-clones. These can be further categorized as early and late subclones based on the overall subclonal architecture of a cancer sample. Thus, we explored the relative population of high- and low-impact putative passengers in different sub-clones of a tumor sample^11^ to decipher their progression during tumor evolution. Intuitively, one might hypothesize that high-impact mutations achieve greater prevalence in tumor cells if they are advantageous to the tumor, and a lower prevalence if deleterious. As expected, we observed this to be true among driver variants (**Fig 4a**). However, interestingly, we observed that high-impact putative passengers in coding regions have greater prevalence among parental subclones (**Fig 4a**) – an effect driven by high-impact putative passenger SNVs in tumor suppressor and apoptotic genes (**Fig 4a**). In contrast, high-impact putative passenger SNVs in oncogenes appear slightly depleted in parental subclones. Similarly, high-impact putative passengers in DNA repair genes and cell cycle genes are depleted in early subclones (**Fig 4a**). We obtained similar results when we simply categorized mutations on the basis of variant allele frequency (VAF) (**supplement Fig. S19**). We note that prior analysis^11^ suggests little difference in signatures between early and late subclone mutations. Thus, signatures are unlikely to drive the observed impact score variations^11^ between early and late subclone mutations.

**Figure 4:**
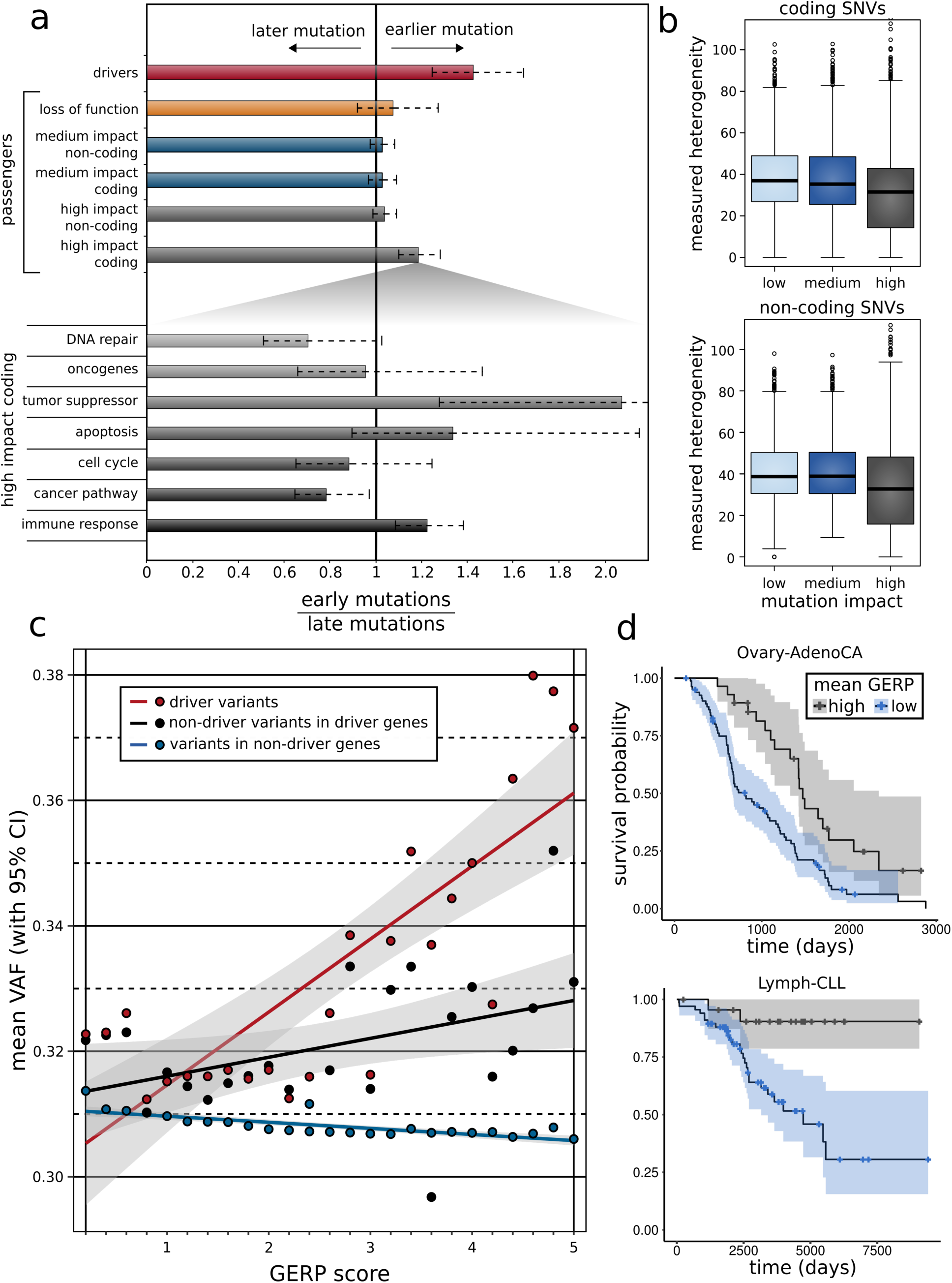
Correlating functional burdening with subclonal information and patient survival. **a)** Subclonal ratio (early/late) for different categories of SNVs (coding/non-coding) based on their impact scores. Subclonal ratios for high-impact SNVs occupying distinct gene sets; **b)** Mutant tumor allele heterogeneity difference comparison between high-, medium-, and low-impact SNVs for coding(top) and non-coding regions(bottom); **c)** Correlation between mean VAF and GERP score of different categories of variants (driver SNVs, non-driver SNVs in known cancer genes, and putative passenger variants in non-driver genes) on a pan-cancer level;**d)** Survival curves in CLL (*left panel*) and RCC (*right panel*) with 95% confidence intervals, stratified by mean GERP score.

In non-rearranged genomic intervals, the VAF of a mutation is expected to be proportional to the fraction of tumor cells bearing that mutation. Previous studies^19^ have measured the divergence in VAFs to indirectly quantify heterogeneity in mutational burden among different sub-clones in a cancer. Here, we quantified this heterogeneity among low-, medium-, and high-impact putative passengers for different cancer cohorts (see **supplement section 6.2)**. We generally observed lower mutational heterogeneity among high-impact putative passenger SNVs. This observation is consistent for both coding and non-coding putative passenger variants (**Fig 4b**).

Furthermore, we correlated the predicted molecular functional impact (measured by GERP score here) of each variant with their corresponding cellular prevalence measured by VAF (see **supplement section 6.3)**. We find that, within driver genes and their regulators, variants that disrupt more conserved positions (high GERP score) tend to have higher VAF values (**Fig 4c**). This trend remains true even after excluding SNVs that have been individually classified as driver variants, suggesting that within driver genes, yet-uncalled driver variants remains (potentially among high impact putative passengers). We also find that outside of driver genes, variants that disrupt more conserved positions tend to have lower VAF values (**Supplement Table 1**).

As with the clonal status of a tumor, clinical outcomes (such as patient survival) provide an alternative measure of tumor progression. Therefore, we performed survival analysis to see if somatic molecular impact burden – here measured as the mean GERP of putative passenger mutations per patient – predicted patient survival within individual cancer subtypes. Patient age at diagnosis was used as a covariate in the survival analysis. We obtained significant correlations between somatic molecular impact burden and patient survival in two cancer subtypes after multiple test correction (**Supplement Table 2**). Specifically, we observed that somatic molecular impact burden predicted substantially better patient survival in lymphocytic leukemia (Lymph-CLL, p-value 2.3e-4) and ovary adenocarcinoma (Ovary-AdenoCA, p-value 2e-3) compared to other cancer types (**Fig 4d**). The use of *average* impact rather than summed impact ensures that these results do not simply reflect more advanced progression (i.e. more mutations) of the cancer at the time of sequencing. We note that unmeasured patient clinical characteristics or tumor molecular subtypes may partially influence these correlations.

## Categorizing putative passenger variants

The comprehensive characterization of the putative passenger landscape in PCAWG highlights many key attributes of putative passengers. The results we have found may be explained in relation to underlying mutational processes. However, they may also be indicative of selective effects among a subset of these mutations, whether or not they are generated by a neutral mutational process. If a subset of putative passengers indeed possesses fitness effects, then we can extend the canonical model of drivers and passengers into a continuum model. Conceptually, in such an extended model, somatic variants can be classified into multiple categories while considering their impact on tumor cell fitness: drivers with strong positive selective effects, and putative passengers with neutral, weak positive, and weak negative selective effects. This broad classification scheme can be further refined by considering ascertainment-bias and the putative molecular functional impact of different variants (**Fig 5a**). Previous power analyses^20,21^ suggest that existing cohort sizes support the identification of strong positively-selected driver variants, but that many weaker drivers and even some moderately strong driver variants would be missed.

**Figure 5.**
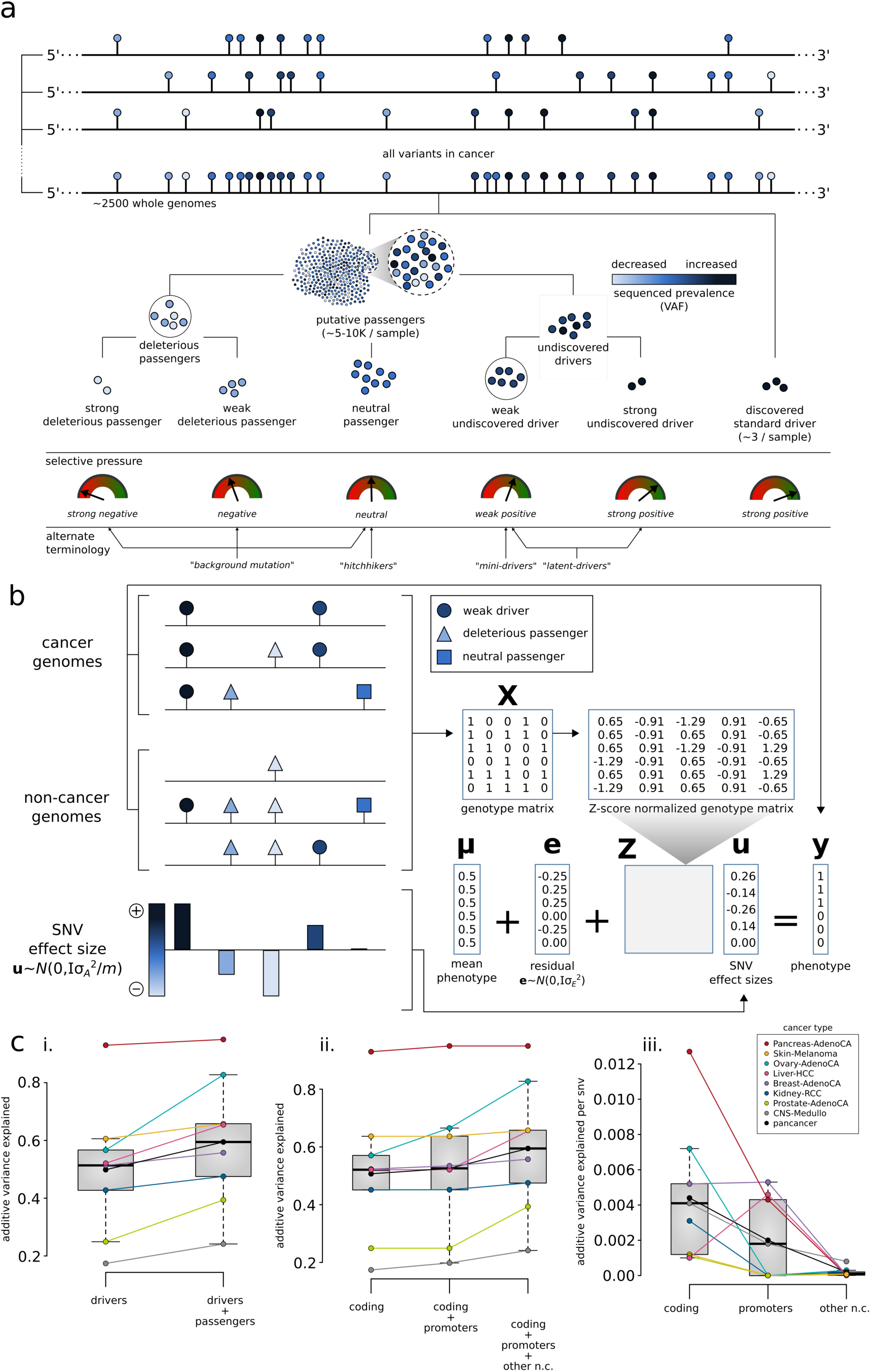
Conceptual classification of somatic variants based on their functional impact and selection characteristics, and additive effects model. **a)** Both coding and non-coding variants can be classified as drivers or putative passengers based on their impact and signatures of positive selection. Among putative passengers, true passengers undergo neutral selection and tend to confer low functional impact. Deleterious passengers (weak and strong) and mini-drivers (weak and strong) represent various categories of higher-impact nominal passenger variants, which may undergo weak negative or positive section; **b)** *Additive effects model for nominal passengers*: The combined effect of many nominal passengers are modeled using a linear model, which predicts whether a genotype arises from an observed cancer sample or from a null (neutral) model (notation defined in **supplement section 8.1**). The model is fitted by optimizing the hyper-parameter 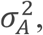, and a test for significant combined effects of the nominal passengers is made by performing a log-likelihood ratio test against a restricted model which includes only µ and e; **c)** *Predictive power of known drivers and nominal passengers using the additive effects model*: (i) compares the maximum possible variance which can be explained using known drivers with the performance of the model containing driver mutations and putative passengers;(ii) further the variance-explained into a contribution from coding variants, non-coding and promoter variants, and all variants; (iii) presents normalized additive variance explained exclusively by putative passengers in coding regions, exclusively by promoters, and exclusively by other non-coding elements of the genome.

However, these moderately strong and weak driver variants can also provide a potential fitness advantage to tumor cells. With respect to a functional-impact-based classification, any positively or negatively selected variants will have some molecular functional impact (i.e. effect on gene expression or activity). The relevance of molecular functional impact is firmly established for driver mutations, defined as positively-selected variants promoting tumor growth. However, a rapid accumulation of *putative passengers*, which undergo weak/strong negative selection, could adversely affect the fitness of tumor cells^6^. Moreover, a majority of low-impact and some high-impact *putative passengers* may alter tumor gene expression or activity in ways that are not ultimately relevant for tumor fitness; hence, these variants will undergo neutral evolution.

An initial step towards identifying the presence of variants with effects on tumor fitness is to compare observed mutation distributions with ones generated by simulating or modeling neutral processes. This approach has been extensively leveraged in the context of individual driver discovery using element burden testing. Such an approach is potentially powerful since it allows the use of complex background mutational models. However, the possibility of detecting artifacts due to the inadequacy of current models of neutral mutational processes remains, since unmodeled mutation processes may result in confounding effects. With this caveat, we explore such an approach below in an attempt to quantify non-neutral aggregate effects among putative passengers, using a variety of background models and an additive effect model which combines both positive and negative fitness effects. As in the case of individual driver discovery, validation of such effects requires follow-up experimentation.

## Overall effects of putative passengers and additive variance

It is interesting to note that in a cancer genome, the presence of few drivers (with high positive fitness effects) and large numbers of putative passengers (with weak or neutral fitness effects) could be considered analogous to prior observations in genome-wide association studies (GWAS) that implicated a handful of variants influencing complex traits. These modest numbers of variants explain only a small proportion of the genetic variance, thus contributing to the “missing heritability” problem in GWAS^22,23^. However, it has been shown that aggregating the remaining variants with weak effects can explain a significant part of the “missing heritability”^22^ and is predictive of phenotype^24^. We do not currently have estimates of “missing heritability” at the subclone level for tumorigenicity (see **supplemental section 8**), which may depend on both genetic and epigenetic factors. However, the fact that some tumors in PCAWG (∼10% of PCAWG samples) lack a known driver^12^ suggests that some driver mutations remain to be discovered. The models above suggest the importance of investigating the cumulative effect of putative passengers in this context.

To address this, we adapted an additive effects model^22,25^, originally used in complex trait analysis, to quantify the relative size of the aggregated effect of putative passengers in relation to known drivers. With a number of caveats regarding interpretation arising due to differences between germline and cancer evolutionary processes (see **supplemental section 8**), we tested the ability of this model to predict cancerous from null samples as a binary phenotypic trait (**Fig 5b**). Briefly, we created a balanced dataset of observed tumor and matched neutral (null) model samples, using a recently proposed background model which preserves mutational signatures, local mutation rates, and coverage bias (see **supplemental section 1.1.b**). Subsequently, using a linear model, the additive effects model implicitly associates a positive or negative effect (coefficient) with each SNV, considering them to be sampled from a normal distribution (see **supplemental section 8.1**). Furthermore, in this model the individual effects of SNVs are not explicitly estimated; instead, their variance is evaluated as a hyper-parameter using restricted maximum-likelihood (REML)^25^, where separate variance terms can be associated with different groups of SNVs falling in distinct categories. In addition to the neutral model above, we further utilized two additional local background models, including PCAWG-wide randomized datasets as well as our custom randomization correcting for various covariates (see **supplemental section 1.1.a-c**).

We compared several versions of the additive variance model (explained above) in 8 cancer cohorts having a sample size greater than 100. In the first model, we separated the mutations into two categories, corresponding to drivers (from the PCAWG analysis) and putative passengers (**Fig. 5ci**). Putative passengers were only included in the model if found in at least two samples from a cohort (which can be any combination of observed and simulated samples). Additionally, to maximize the predictive potential of the driver mutations, we used a binary variable which is 1 if any driver mutation is present in a sample as a predictor (details in **supplement section 8.1**). This approach effectively isolates the effect of putative passengers in tumors without driver mutations. In this model, we observed an increase in the variance explained from ∼49.9% using drivers alone to ∼59.4% with putative passengers when averaged across all cohorts, with the putative passenger contribution significant at FDR<0.1 in all cohorts except kidney-RCC cohort, suggesting that non-neutral effects are present among the putative passenger mutations (**Supplement Table 3a**). We further tested a different version of the model in which we split mutations into coding, promoter, and other non-coding categories, where the coding mutations are a superset of the PCAWG drivers (**Fig. 5cii**). Here, we observed that the coding mutations by far accounted for the largest overall proportion of the variance (∼50.7% averaged across cohorts), while promoter and other non-coding mutations contributed much less, but still significant amounts of extra variance (∼1.9% and 6.9% respectively overall, with cohort-specific contributions from each category at FDR<0.1, **Supplement Table 3b**). Although the total contribution of the promoters is lowest in this model, the additive variance per SNV (normalized variance) is substantially higher in promoters than other non-coding mutations (**Fig. 5ciii**). However, the normalized additive variance for promoter is lower than coding mutations. Further, we tested the sensitivity of our results to the choice of null model by repeating these analyses for two other randomization schemes, and compared the variance on observed and liability scales, with quantitatively similar results (**Supplement Tables 4-7**). We also verified that our model effectively controls for overfitting by observing near zero additive variance when a second randomized sample is substituted for the observed genotypes, and that the sensitivity of the results to changes in the randomization window size is small (**Supplemental Table 10a-b**). To further verify that the model controls for overfitting, we split the data into test and training partitions, and showed that the additive variance on the training partition correlates with test predictive accuracy, where we use the Best Linear Unbiased Predictor (BLUP) to cast the model in predictive form (**Supplemental Table 10c and supplemental section 8.2**).

By including a binary predictor for known driver SNVs in the above model, we expect the contribution of the putative passengers to be higher among samples without known drivers (as well as all null samples). To confirm that the putative passengers were indeed contributing to the discrimination of samples without known drivers, we further calculated the additive variance exclusively for such samples in PCAWG. First, we repeated the analysis of the 8 cohorts above, but with all samples containing known SNV drivers and SNVs falling in known driver elements removed. We observed an average of 12.5% additive variance across cohorts (**Supplement Table 8a**), which was higher than the 9.5% additive variance estimates based on putative passengers among all samples (with and without known drivers; p=0.01, 1-tailed paired t-test for increase in per-cohort additive variance, all cohorts ≥ 20 samples). This observation is consistent with a more important role for the putative passengers among samples without a known driver, since they may have partially redundant effects in the samples harboring known drivers, or include a larger number of high-impact undiscovered drivers among the samples without known drivers. In a subsequent analysis, we calculated the additive variance after excluding samples with driver SVs and CNAs in addition to samples with known driver SNVs. This analysis was performed for pan-cancer meta-cohort which pools all such samples (**Supplemental Table 8b**). We observed a lower amount of additive variance (6.8%) for the pan-cancer meta cohort, which may be due to tissue-specific effects which are lost at the meta-cohort level. Further, we tested for possible inflation of BLUP coefficients on q arm of the chromosome1(1q) due to gain of 1q events in both the analyses above, but found no effect across cancer cohorts (**Supplemental Table 11)**, suggesting that arm-level aneuploidy does not contribute appreciably to these estimates.

Finally, we estimated the BLUP for individual cohorts after samples with predicted SNV, SV and CNA drivers were excluded, including SNVs contained in predicted driver elements (details in **supplement section 8.2**), and used this to derive an estimate of the number of weak drivers among samples lacking predicted PCAWG drivers^12^ (**Supplement Table 9**). We conservatively estimated the number of weak drivers by finding the smallest set of SNVs whose additive variance is equal to the additive variance of all the SNVs. A per sample estimate of driver events is then derived by comparing the average number of SNVs from this set in the observed versus random samples. Using this approach, we estimated an average of 8.4 weak drivers per cohort, corresponding to approximately 0.81 weak driver events per tumor. We expect that these estimates are limited by sample size, and thus represent conservative lower bounds.

## Discussion

Certain key alterations in the tumor genome, often identified through the detection of strong signals of positive selection on individual variants, have been shown to play a pivotal role in tumor progression. Although a typical tumor has thousands of genomic variants, very few of these (∼5/tumor^4^) are thought to drive tumor growth. The remaining variants, often termed passengers, represent the overwhelming majority of the variants in cancer genomes, and their functional consequences are poorly understood. In this work, we comprehensively characterized putative passengers in the PCAWG dataset. Subsequently, we attempted to quantify the cumulative fitness effect of such putative passengers on tumor growth through the additive variance model. We note that the above approach relies on applying an accurate background model. However, current null models have inaccuracies due to our incomplete understanding of various mutational processes in cancer. Nonetheless, our additive variance analysis was robust for multiple background models and suggested a potential for putative passengers to play a role in tumor progression through their cumulative effect. This effort further complements the driver discovery exercise^9^ in PCAWG by identifying key alterations beyond strong drivers. Also, our predicted functional impact analyses of putative passengers showed that different mutational processes are associated with extensive differences in impact on cellular subsystems, irrespective of whether these cause, are indirectly associated with, or are independent of subclonal fitness differences in an evolving tumor. These observations further motivate follow-up experiments and additional whole-genome analyses to explore the role of putative passengers with weak (positive and negative) fitness effects in cancer. In conclusion, our work highlights that an important subset of somatic variants currently identified as putative passengers nonetheless may have biologically and clinically relevant functional roles across a range of cancers.

## Acknowledgements

MG acknowledge support from the NIH and from the AL Williams Professorship funds. We are thankful to the members of the PCAWG technical working group for generating the variant calls. We are also thankful to the members of the PCAWG steering committee for providing valuable feedback on the manuscript.

